# Continuous exposure to 60 Hz extremely low frequency electromagnetic field at 10 to 16 mT promotes various human cell proliferation by activating extracellular-signal-regulated kinase

**DOI:** 10.1101/2024.06.12.598738

**Authors:** Jaeseong Goh, Donghwa Suh, Dae Yong Um, Seung Ahn Chae, Gwan Soo Park, Kiwon Song

## Abstract

We previously showed that continuous exposure to 60 Hz extremely low-frequency electromagnetic fields (ELF-EMF) at 6 mT promotes cell proliferation. Here, we investigated the cellular effect of 60 Hz ELF-EMF at over 10 mT. We revised the ELF-EMF-generating device to increase the magnetic flux density of the ELF-EMF stably without thermal effect. We investigated the cellular effect of 10-16 mT ELF-EMF on various mammalian cells including human cervical carcinoma HeLa, rat neuroblastoma B103, liver cancer stem cells Huh7 and Hep3B, immortalized normal hepatic cell MIHA, and normal fibroblast IMR-90. Cell proliferation was promoted around 20% or more in all cells through continuous ELF-EMF exposure at 10 and 14 mT for 72 h, compared with the sham exposure group. In the cells whose proliferation was activated by 14 mT ELF-EMF, the MEK-ERK pathway and NF-κB were activated but not Akt. These cells showed a slight increase in the S phase population in BrdU incorporation and Ki-67 expression. In these cells, intracellular and mitochondrial ROS levels were not changed, and the proliferation-activating cellular effects of ELF-EMF were maintained even when oxidative phosphorylation was interrupted by CCCP. Additionally, no changes in intracellular calcium levels were observed in ELF-EMF-exposed cells and the proliferation-activating cellular effects of ELF-EMF were maintained in the presence of a calcium chelator, BAPTA-AM. These observations suggested that ROS and intracellular calcium do not mediate ELF-EMF’s proliferation-activating physiological effect. Altogether, we demonstrated that 60 Hz ELF-EMF at 10 to 14 mT promotes cell proliferation by activating ERK1/2 and does not affect intracellular ROS and calcium levels.

## Introduction

Extremely low-frequency electromagnetic fields (ELF-EMFs) are a type of non-ionizing radiation with frequencies ranging from 3 to 3000 Hz [1, 2]. ELF-EMFs are ubiquitous in our surroundings, predominantly originating from power lines, electrical appliances, and industrial processes. With rapid technological advances, the growing number of sources emitting ELF-EMFs is inevitable [3]. Given the expansion of ELF-EMF exposure in daily environments, the safety of ELF-EMF exposure has been one of the major public concerns for decades, and scientific evidence is needed to ensure protection from its potential adverse effects. Since the 1980s, numerous epidemiological studies have indicated potential associations between ELF-EMFs and increased incidence of leukemia, lymphoma, and nervous system tumors [4–8]. However, the findings of these studies were frequently obscured by ambiguity because of methodological weaknesses, and incomplete variable control [9].

Several *in vitro* studies have claimed that ELF-EMFs positively influence various stages of the complex wound healing process, including inflammation modulation, tissue rearrangement, and cell proliferation, with effects varying depending on exposure time, waveform, frequency, and amplitude [10]. A quite number of studies have explained the physiological changes in cells caused by ELF-EMF exposure to the generation of reactive oxygen species and the influx of calcium ions [11, 12]. In particular, exposure to 50 Hz ELF-EMF at 1 mT induced neural differentiation in human bone marrow mesenchymal stem cells (BM-MSCs) via EGFR signaling activation and mild ROS increase [13]. Exposure to 50 Hz, 1 mT ELF-EMF for 6 h daily enhanced cell proliferation, de novo calcium deposition, and osteogenic differentiation in human periodontal ligament mesenchymal stem cells [14]. ELF-EMF exposure at 50 Hz, 1 mT for 4 h per day enhanced embryonic neural stem cell (eNSC) proliferation and neuronal differentiation, and increased calcium levels, linked to elevated TRPC1 expression [15]. Some other studies also demonstrated that ELF-EMF activates various stress response signaling pathways. Patruno *et al.* reported that exposure to 50 Hz ELF-EMF at 1 mT promoted the proliferation of HaCaT cells by activating PI3K/AKT and ERK pathways [16]. Exposure to 50 Hz ELF-EMF at 1 mT promoted cellular growth through the ROS-mediated NF-κB pathway in HL-60, Rat-1, and WI-38 cells [17].

While most 50 ∼ 60 Hz ELF-EMF studies were conducted under 1 mT, we wanted to examine the effect of ELF-EMF at over 10 mT, which occurs around social infrastructure such as large power plants, factories, and telecommunication facilities. We previously reported that uniform 60 Hz ELF-EMF at under 6 mT shows neither genotoxic nor apoptotic effects, and rather promotes cell proliferation in both cancer and normal human cells [18]. In this study, we designed a novel device to generate uniform 60 Hz ELF-EMF at 10 to 20 mT without generating heat. With this device, we investigated the cellular effects of 60 Hz ELF-EMF at over 10 mT on various human cells and tried to elucidate the factors responsible for the physiological responses.

## Materials and Methods

### Sources of cells and culture

Human adenocarcinoma HeLa cells and human lung fibroblast IMR-90 cells were purchased from the American Type Culture Collection (ATCC), and human hepatocellular carcinoma Huh7 cells were purchased from the Korean Cell Line Bank (KCLB). Human hepatocellular carcinoma Hep3B cells were provided by Dr. Y.N. Park (Yonsei University College of Medicine, Seoul, Korea). B103 rat neuroblastoma cells were a gift from Dr. Inhee Mook-Jung (Seoul National University College of Medicine, Seoul, Korea). Immortalized normal hepatic cell MIHA was a gift from Dr. S.W. Nam (The Catholic University College of Medicine, Seoul, Korea).

HeLa, B103, Huh7, Hep3B, and MIHA cells were cultured in high glucose-containing Dulbecco’s modified Eagle’s medium (DMEM; Gibco, USA, #11965092) supplemented with 10% fetal bovine serum (FBS; Sigma-Aldrich, St. Louis, MO, USA, #26140-079) and 1% penicillin-streptomycin (Gibco, USA, #2585855). IMR-90 cells were grown in a minimum essential medium (MEM; Gibco, USA, #11095080) supplemented with 10% FBS and 1% penicillin-streptomycin. All cells were cultured at 37℃ in a humidified atmosphere containing 5% CO_2_.

### Exposure of cells to 60 Hz Extremely low frequency-electromagnetic field

The same number of cells (3 × 10^4^) was seeded in 35 mm cell culture dishes and incubated for 18 h for both the ELF-EMF-exposed group and the sham exposure group. Then, the cells of the exposed group were moved to the ELF-EMF device to be exposed to a 60 Hz uniform ELF-EMF at the indicated intensity and durations, while the sham exposure group was incubated during the same durations as the exposed group in the same incubator. At least three sets of cells in separate dishes with three sets of sham exposure groups were exposed in each exposure experiment. A customized thermometer was used to ensure the temperature was at 37 ± 0.5°C, both inside the ELF-EMF device and in the incubator.

### Cell viability assay

Cell viability was monitored using MTT assays. For both the ELF-EMF-exposed and the sham exposure groups, cells were incubated in 1 mL of medium containing 0.5% 3-(4,5-dimethylthiazol-2-yl)-2,5-di-phenyltetrazolium bromide (MTT; Amresco Inc., OH, USA, #97062-380) at 37℃ for 90 min. The resulting formazan crystals were dissolved in 1 mL of dimethyl sulfoxide (DMSO, Duksan, Korea, #1380), and the optical density was measured at 570 nm using an ELISA microplate reader (SpectraMax ABS, Molecular Device Co., CA, USA).

### Western blot analysis

Cells were harvested and washed with cold phosphate-buffered saline (PBS). The collected cells were lysed with lysis buffer containing 50X Protease Inhibitor Complex (PIC; Sigma-Aldrich, Germany, #11836170001) and 100 mM phenylmethylsulfonyl fluoride (PMSF, GaBiochem, Inc., USA, #329-88-6) and the protein concentration of each lysate was measured with Bicinchoninic acid assay (BCA assay, Thermo Scientific, MA, USA, #23328 and #23224) as previously described [18]. Each lysate containing the same amount of proteins was separated on 8–12% SDS-polyacrylamide gels (PAGE) and electro-transferred to PVDF membranes (Merck Millipore, Billerica, MA, USA, #IPVH00010). The membrane was incubated at 4°C overnight with the following primary antibodies: anti-phospho-MEK1/2 (Cell Signaling Technology, Inc., MA, USA, #9122), anti-MEK1/2 (Santa Cruz biotechnology, INC., #SC-7995-R), anti-phospho-ERK1/2 (Santa Cruz biotechnology, INC., #SC-7383), anti-ERK1/2 (Cell Signaling Technology, Inc., MA, USA, #9102), anti-phospho-Akt (Cell Signaling Technology, Inc., MA, USA, #9271), anti-Akt (Cell Signaling Technology, Inc., MA, USA, #9272), anti-phospho-p65 (Cell Signaling Technology, Inc., MA, USA, #3033), anti-p65 (Cell Signaling Technology, Inc., MA, USA, #8242), anti-Cdk4, anti-PCNA (Santa Cruz Biotechnology, TX, USA, #SC-56). After washing with PBS (phosphate-buffered saline), the membrane was incubated with anti-mouse (Cell Signaling Technology, Inc., A, USA, #7076) or anti-rabbit (Cell Signaling Technology, Inc., A, USA, #7074) secondary antibody at room temperature for 1 h. Chemiluminescence was detected with Amersham^TM^ ImageQuant 800^TM^ (Cytiva). All blots were stripped and re-probed with anti-ꞵ-actin (Cell Signaling Technology, Inc., #4970) for loading controls. The band intensity relative to the loading control (ꞵ-actin) was measured using ImageJ software (NIH, MD, USA).

### Cell cycle and cell death analysis

For cell cycle analysis, ELF-EMF-exposed cells were processed as described [18] and stained with 50 μg/mL propidium iodide for 1 h at 4℃ in the dark. Cells were also processed with FITC BrdU Flow Kit (BD Pharmingen™, USA, #559619) for BrdU-7AAD staining, following the manufacturer’s instructions. The distribution of cells in the cell cycle and the BrdU incorporation were analyzed by flow cytometry. For cell death analysis, cells resuspended in 1X Annexin V binding buffer were incubated with FITC conjugated Annexin V and propidium iodide (PI; BD Pharmagen^TM^) and analyzed by flow cytometry. Flow cytometry was performed with Accuri™ C6 Plus (BD Bioscience, NJ, USA), and 10,000 cells per assay were analyzed with FlowJo V10 software (Tree Star, CA, USA).

### ROS measurement

Intracellular and mitochondrial ROS levels were measured using a ROS detection kit (Invitrogen, MA, USA, #88-5930-74) and Mitosox Red (Invitrogen, MA, USA, #M36008), respectively, following the manufacturer’s protocol. The ELF-exposed and sham control cells were washed with PBS and incubated with 25 μM carboxy-H_2_DCFDA for 30 min or 5 μM MitoSox for 10 min at 37°C in the dark. After incubation, cells were harvested by trypsinization and analyzed by flow cytometry with BD Accuri™ C6 Plus (BD Bioscience, NJ, USA). 10,000 cells per assay were analyzed with FlowJo V10 software (Tree Star, CA, USA).

### Intracellular calcium measurement

Intracellular calcium levels were evaluated with Fluo-4-AM (Invitrogen, MA, USA, #F14201). ELF-EMF-exposed cells were incubated with 1 ml of medium containing 2.5 μM Fluo-4-AM for 30 min. After incubation, cells were harvested and analyzed with BD Accuri™ C6 Plus. The mean fluorescence intensity was calculated by FlowJo V10 software (Tree Star, CA, USA).

### Immunofluorescence Microscopy

Cells were seeded onto collagen-coated coverslips (25 mm) in 35 mm dishes, fixed in 100% methanol, and then permeabilized with 0.5% Triton X-100. After permeabilization, the cells were blocked with 3% BSA for 1 h at room temperature, followed by immunostaining for Ki-67 and mounted with 4,6-diamidino-2-phenylindole (DAPI). An anti-Ki-67 antibody (Cell Signaling Technology, Inc., USA, #15580) was used as the primary antibody, and Alexa Fluor 488 goat anti-rabbit IgG (Invitrogen, MA, USA, #A11008) was used as the secondary antibody. The cells were visualized on a Zeiss Axioplan2 fluorescence microscope (Carl Zeiss, Gina, Germany), and images were acquired using an Axiocam CCD (Carl Zeiss, Gina, Germany) camera and AxioVision software (Carl Zeiss, Gina, Germany).

### Transcriptome analysis

B103 cells treated with 60 Hz ELF-EMF at 14 mT for 24 h were harvested for RNA extraction and whole genomic RNA-sequencing was performed by Macrogen Co. (Republic of Korea). Differentially expressed genes (DEGs) were identified using edgeR between the ELF-EMF exposed group and the unexposed control. Genes with a p-value of less than 0.05 and more than a log2 fold change of 1.3 were considered to be DEGs. Functional enrichment analysis was performed with DEGs using the Database for annotation, visualization, and integrated discovery (DAVID).

### Chemical treatment

After 18 h of cell seeding, the cell culture medium was removed and replenished with a culture medium respectively containing carbonyl cyanide 3-chlorophenylhydrazone (CCCP; Sigma-Aldrich, MA, USA, #C2759), and 1,2-bis(2-aminophenoxy)ethane-N,N,N′,N′-tetraacetic acid acetoxymethyl ester (BAPTA-AM; MedChemExpress, #HY-100545). The chemicals were dissolved in dimethyl sulfoxide (DMSO) and the same volume of DMSO was added to the untreated controls.

### Statistical analysis

All statistical analyses were performed using GraphPad Prism 9 statistics program (GraphPad Software Inc., San Diego, CA, USA). All data are presented as mean ± standard deviation (SD) of more than three independent experiments with statistical significance. P<0.05 (*), and P<0.01 (**) were considered to indicate statistical significance, while P>0.05 was considered to indicate statistical non-significance (ns).

## Results

### An ELF-EMF generating device was improved to apply a magnetic flux density of 10-20 mT

In the previous study, we examined the biological effect of 60 Hz EMF exposure at 6 mT on human cells with a closed-type EMF device that generates a uniform ELF-EMF [18]. However, when the magnetic flux density was increased to 10 mT with this device, heat was generated. Because heat also affects the physiology of cells, we could not use this device to investigate the biological effect of 60 Hz ELF-EMF at over 6 mT magnetic flux density. In this study, in order to examine the effect of 60 Hz ELF-EMF on human cells at 10 mT or higher magnetic flux density, we re-designed and manufactured an improved ELF-EMF device, in which the magnetic flux density can be increased up to 20 mT without a heat-generating problem.

The length of the device is 280 mm, the number of turns is 1500, and the wire diameter is 1 mm. The resistance and inductance of the prototype device are 26.02 Ω and 3.26 H, respectively. Thus, the device generates 0.087 mT per unit supply voltage, so that up to 20 mT can be achieved at a supply voltage of 230 V. The specification of the device is summarized in Table 1. The schematic pictures of the previous and the newly revised devices are shown in Supplementary Figure S1.

**Table 1.**
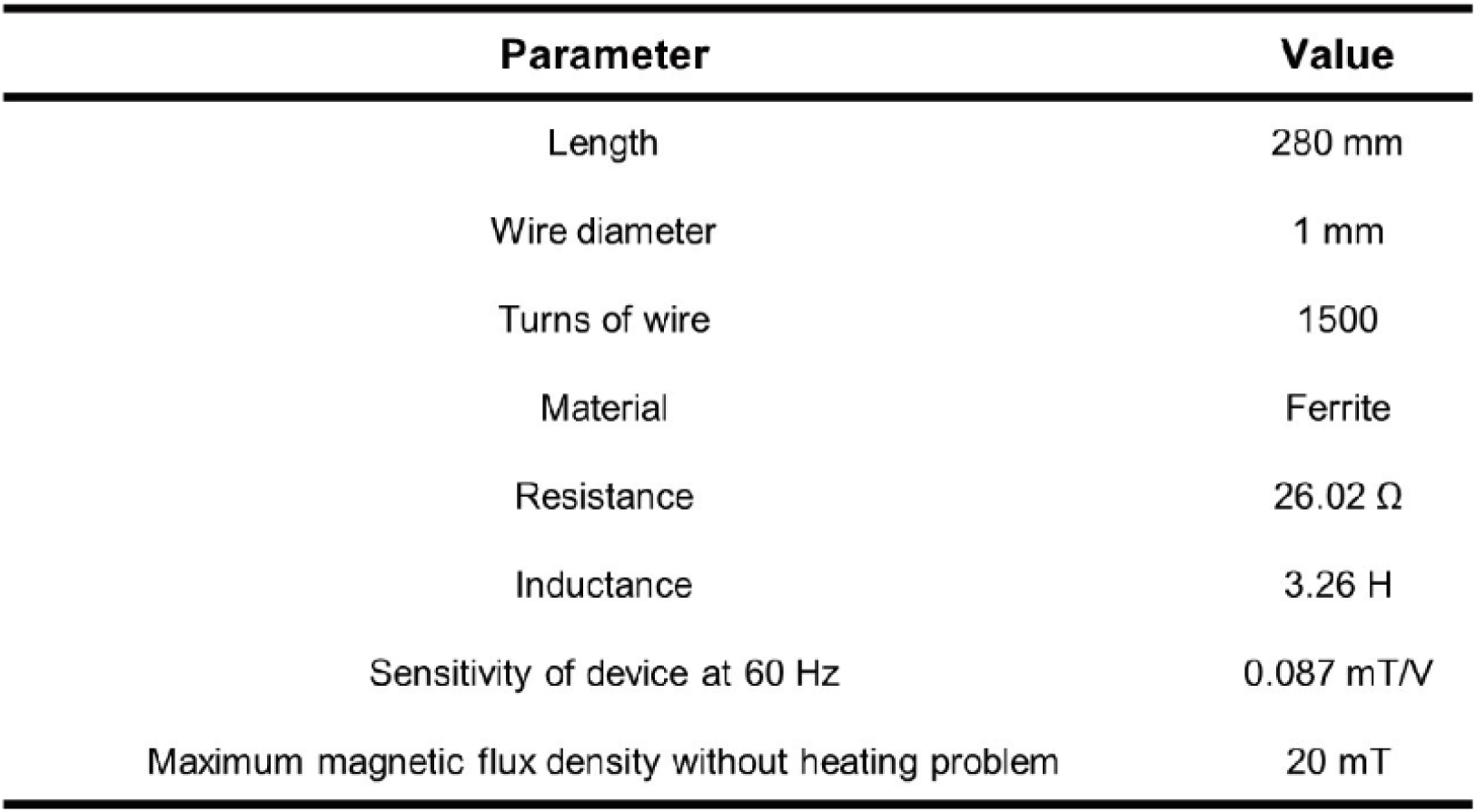
Design parameters for the ELF-EMF generating device.

| Parameter | Value |
| --- | --- |
| Length | 280 mm |
| Wire diameter | 1 mm |
| Turns of wire | 1500 |
| Material | Ferrite |
| Resistance | 26.02 $\Omega$ |
| Inductance | 3.26 H |
| Sensitivity of device at 60 Hz | 0.087 mT/V |
| Maximum magnetic flux density without heating problem | 20 mT |

This revised electromagnetic device was designed to apply a strong and uniform magnetic field to cell dishes while suppressing temperature increases in an incubator. The overheating problem is caused by the power dissipation of the coil in the device, which is called ‘Joule’s heating law’. The joule law can be written as:

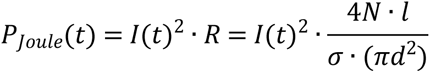

where I is the exciting current into each wire, R is the resistance of the coil, σ is the electrical conductivity of the wire, N is the number of turns, l is the average length per turn of the coil, and d is the wire diameter. To suppress the Joule loss, the exciting current or resistance of the coil should be reduced significantly. Assuming that a high permeability magnetic core is used and a fringing magnetic field around the airgap is neglected, the magnetic flux density on the cell dish can be derived as follows,

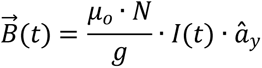

The design of the ELF-EMF generating device is illustrated in Figure 1 A and B. Two experiments with the prototype device were implemented to verify whether the device can expose the uniform magnetic field on each dish and operate properly without overheating. The magnetic field was measured by a Gauss meter (7010; F.W. Bell, Milwaukie, OR, USA) and compared with the simulated result. Both simulated and measured results show their consistency and the magnetic field is almost uniform throughout the cell dish, with the variation of the magnetic field being only 0.3 mT (Fig. 1C). In addition, the temperature variation inside the device was measured. Under operating conditions of 20 mT magnetic field, the device was totally enclosed and operated for a long term while keeping the external temperature at room temperature. The temperature variation was also predicted by the heat-transfer simulation and all parts of the device were modeled (Fig. 1D). The material properties of each part are written in Table 2. The temperature inside the device reaches a steady state after 6 hours (Fig. 1E). There was a slight difference between the experimental and simulation results. Some factors may cause a difference in that the actual experimental condition was not an ideal isothermal condition as in the simulation and airflow inside the device was not reflected in the simulation. However, both results can represent that the temperature variation is within 0.5°C and the device can generate a strong magnetic field without overheating.

**Figure 1.**
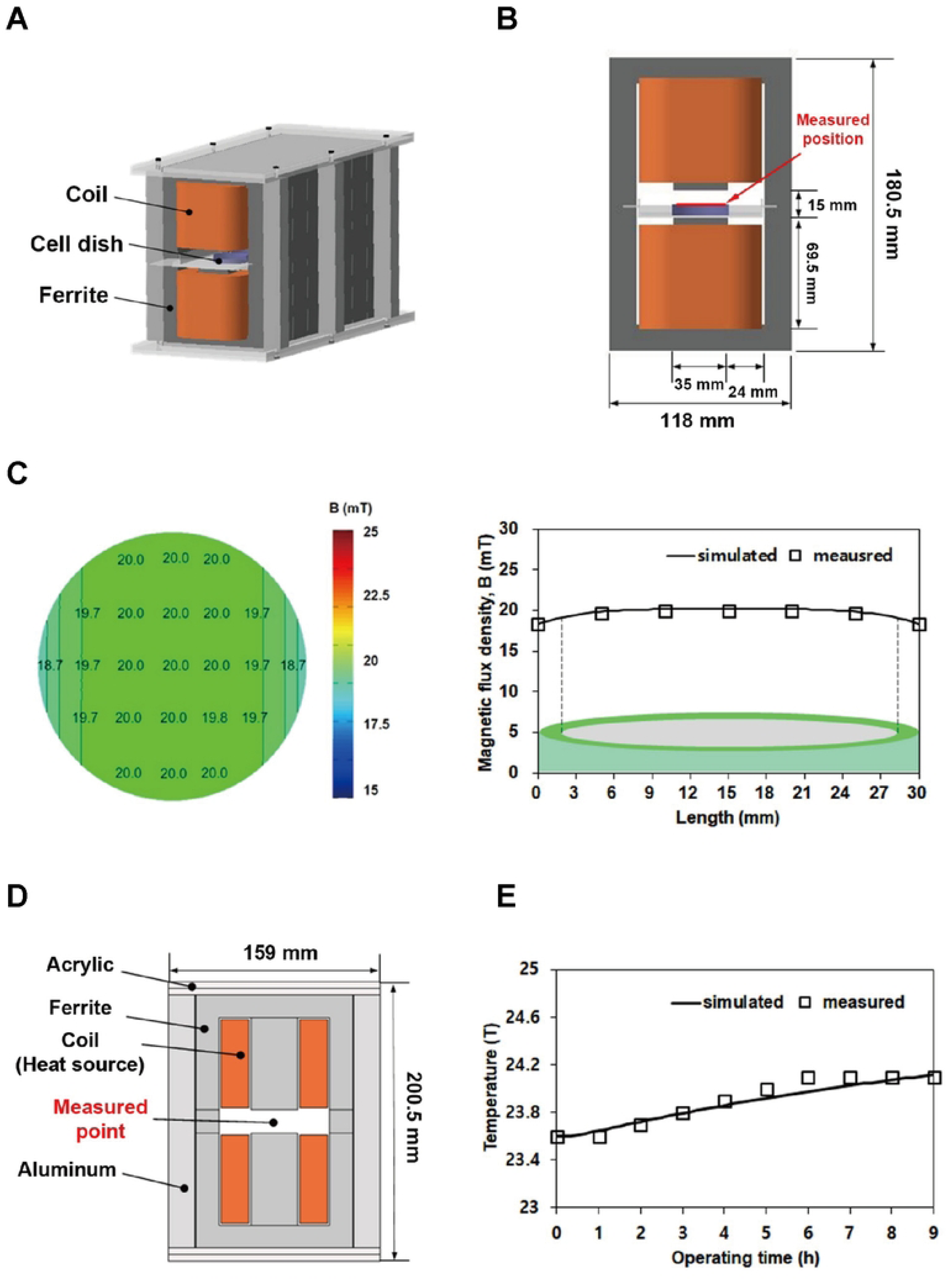
A closed-type device that generates a uniform 60 Hz EMF with 10-20 mT magnetic flux density. (A) An overview of the ELF-EMF-generating device. (B) Schematic front view and size of the device. (C) Simulated and measured magnetic flux density distribution using commercial software, Ansys Maxwell 3D, and a Gauss meter throughout the cell dish. (D) All parts of the device. (E) Simulated and measured temperature changes by using commercial software, COMSOL Multiphysics, and a temperature sensor inside the device.

**Table 2.** Material properties of ELF-EMF generating device.

| Parameter | Air | Ferrite | Copper | Acrylic | Aluminum |
| --- | --- | --- | --- | --- | --- |
| Relative permeability | 1 | 2400 | 1 | 1 | 1 |
| Thermal conductivity [W/(m·K)] | 0.025 | 3.5 | 391 | 0.18 | 210 |
| Heat capacity at constant pressure [J/(kg·K)] | 1007 | 800 | 380 | 1470 | 900 |
| Density [kg/m<sup>3</sup>] | 1204 | 4800 | 8940 | 1190 | 2700 |

To increase the magnetic field on a cell dish, the exciting current should be increased. The heat source that leads to the thermal problem is proportional to the square of the exciting current, thus the overheating issue has arisen in the conventional device. When the voltage is increased, the induced magnetic field rises linearly, but the loss is proportional to the square of the current (I^2^), resulting in overheating. The improved model was designed to avoid the temperature rise in the incubator due to the thermal problem of the device. Compared to the conventional device, the area of the winding window was widened as much as possible to adopt a larger wire diameter within the range maintaining the fundamental role of the ferrite core. Consequently, both the exciting current and the resistance of the coil were optimally minimized so that the source of heating could be extremely suppressed. The overall specifications of both conventional and improved models are shown in Table 3. The total resistance of the improved model is significantly reduced, so that the total heat source of the device decreases from 2.68W to 0.94W, and the thermal problem is alleviated significantly.

**Table 3.** Comparison of previous and improved device.

| Parameter | Previous | Improved |
| --- | --- | --- |
| Length | 280 mm | 280 mm |
| Wire diameter | 0.4 mm | 1 mm |
| Turns of wire | 1000 | 1500 |
| Material | Ferrite | Ferrite |
| Resistance | 104.56 $\Omega$ | 26.02 $\Omega$ |
| Exciting current at 20 mT | 0.16 A | 0.19 A |
| Total heat source of a device | 2.68 W | 0.94 W |

### Continuous exposure to uniform 60 Hz ELF-EMF at 10 to 16 mT promotes cell proliferation

We previously reported that continuous exposure to uniform 60 Hz ELF-EMF at 6 mT increased cell proliferation in HeLa and IMR-90 [18]. In this study, we tried to investigate the physiological effect of 60 Hz ELF-EMF at 10 mT or higher magnetic flux density on various mammalian cells including cancer and normal cells. Before exposing cells to 60 Hz ELF-EMF generated by a newly revised device, we monitored the internal temperature of the device in the incubator at the magnetic flux density of 10, 14, 16, 18, and 20 mT. Notably, at 18 mT and above, a meaningful temperature rise was observed on the inside of the device, compared with that of the outside of the device (Supplementary Figure S2). Thus, in this study, we applied the magnetic flux density within the range that did not significantly increase the internal temperature of the device, 10, 14, and 16 mT.

To examine the effect of 60 Hz ELF-EMF on various mammalian cells, human cervical cancer HeLa, hepatocarcinoma Huh7, hepatocarcinoma Hep3B, lung fibroblast IMR90, immortalized hepatocyte MIHA, and rat neuroblastoma B103 cells were continuously exposed to 60 Hz ELF-EMF at 10, 14, and 16 mT for 72 h. For exposure, at least three sets of the same number of cells plated on 35 mm dishes were pre-incubated for 18 h for both the exposed group and the sham exposure group. Then, the exposed groups were transferred to the ELF-EMF device in the incubator for ELF-EMF exposure at the indicated intensity and durations, while the sham exposure group was incubated during the same durations in the same incubator without ELF-EMF exposure. After 72 h exposure, the cell viability of ELF-EMF-exposed cells was monitored using MTT assays and compared with that of each sham exposure control. As shown in Figure 2, cell viability was promoted in the 60 Hz ELF-EMF-exposed cells compared to the sham controls in all cell types observed and the increased viability was consistent in human and rat cells and in normal and cancer cells. Interestingly, the increase in cell viability was proportional to the increase in magnetic flux density at 10 and 14 mT in all cell types (Fig. 2). Notably we observed the most efficient increase in cell viability at 14 mT in all cell types: the relative increase of viable cells in the exposed over the sham control was 16.9% for HeLa, 25.78% for B103, 29.57% for Huh7, 19.8% for MIHA, 12.37% for IMR 90, and 11.1% for Hep3B. In contrast, at 16 mT, there was no further increase in cell viability as compared to that at 14 mT. Cell viability was even decreased at 16 mT in IMR90 and Hep3B cells when compared to that at 14 mT (Fig. 2 E, F). These results demonstrated that continuous exposure to 60 Hz ELF-EMF at 10 mT to 14 mT consistently increased viable cells in different cell types used, suggesting activated proliferation by the ELF-EMF exposure. They also suggest that the proliferation-activating effect of 60 Hz ELF-EMF depends on the specific magnetic flux density applied.

**Figure 2.**
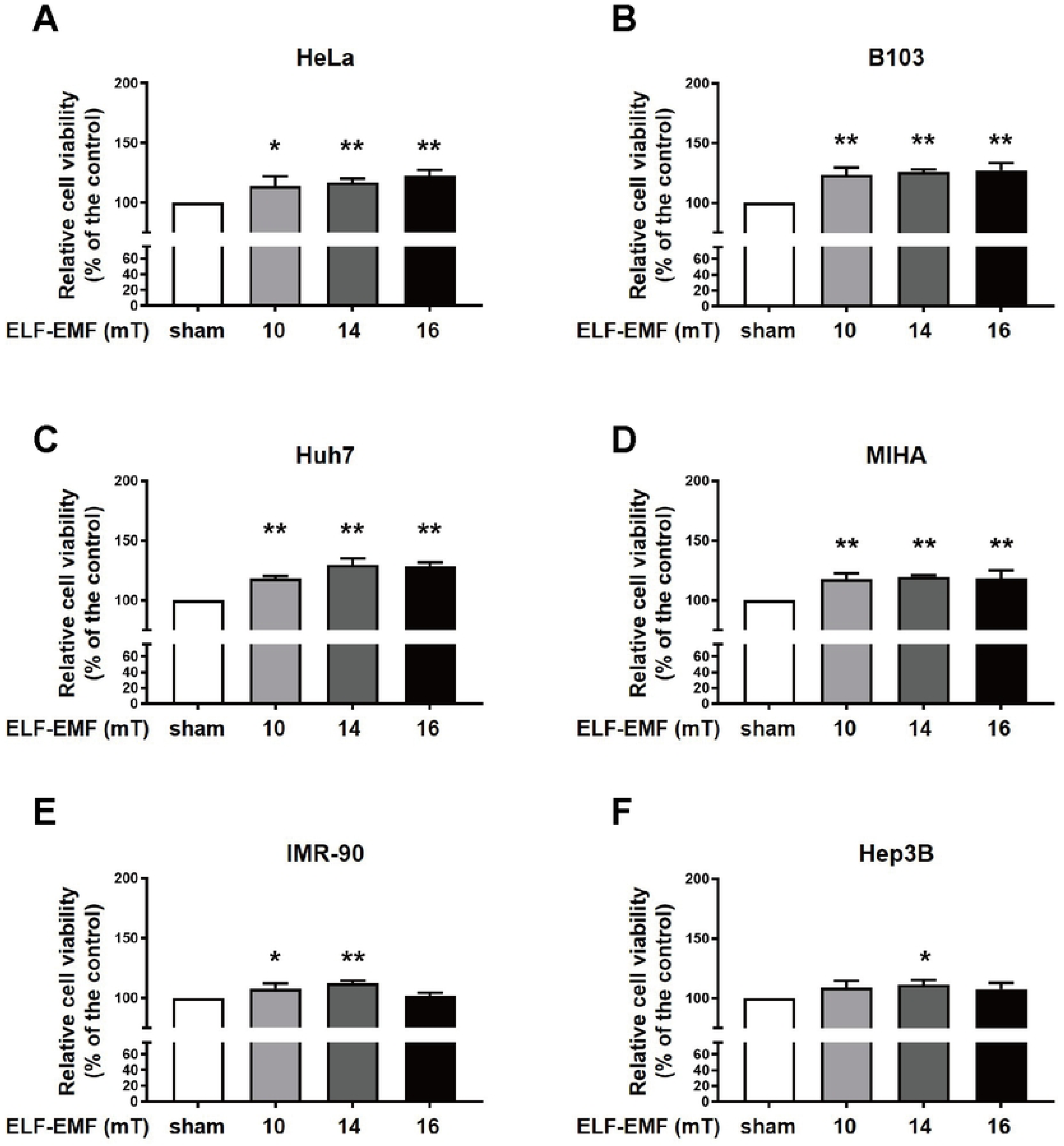
Continuous exposure to 60 Hz ELF-EMF at 10-16 mT for 72 h promoted cell proliferation. (A-F) Human HeLa, Huh7, MIHA, IMR-90, Hep3B, and rat B103 cells were exposed continuously to 10, 14, and 16 mT ELF-EMF for up to 72 h. The sham exposure group of each cell type was incubated for up to 72 h in the same incubator. The cell viability was assessed by MTT assay and the relative cell viability of ELF-EMF-exposed cells was plotted as a percentage of each sham exposure group (100%). Values are presented as the mean ± SD (n = 3). Statistical analyses were performed using GraphPad Prism 9 program (GraphPad Software Inc., San Diego, CA, USA). Data are presented as mean ± standard deviation (SD) of more than three independent experiments with statistical significance. P<0.05 (*), and P<0.01 (**) were considered to indicate statistical significance.

### Continuous exposure to 60 Hz ELF-EMF at 14 mT induces ERK activation

To investigate the mechanism of the cell-proliferative effect induced by 60 Hz ELF-EMF at 14 mT, we examined several signal transduction pathways known to activate the proliferation of cells. Our previous study showed that continuous exposure to 60 Hz ELF-EMF at 6 mT led to increased proliferation via AKT and ERK pathways in HeLa and IMR90 cells [18]. Thus, we first monitored whether the mitogen-activated protein kinase kinase/extracellular-signal-regulated kinase (MEK-ERK) pathway is activated in cells exposed to 60 Hz ELF-EMF at 14 mT. Among different cell types used, human HeLa and rat B103 showed the most highly increased cell viability by 60 Hz ELF-EMF. Thus, we chose HeLa and B103 cells for studying the mechanism of increased viability by ELF-EMF. Considering that cells are most sensitive at the early stage of imposed stress, we examined the activation of MEK-ERK pathway in early exposure durations within the doubling time of HeLa and B103. The doubling time was roughly 18 h for B103 and 25 h for HeLa. HeLa and B103 cells were continuously exposed to ELF-EMF at 14 mT for 6, 12, and 24 h, and ERK phosphorylation was assessed immediately after each exposure. ERK activation by phosphorylation of ERK was detected in both HeLa and B103 cells at all exposure time points assessed, with the most remarkable changes observed at 12 h for HeLa cells and 24 h for B103 cells (Fig. 3A). At the time point that showed the most remarkable ERK activation, we also monitored MEK phosphorylation that activates ERK. Consistently, we observed a slightly increased MEK phosphorylation in 12 h-exposed HeLa cells and in 24 h-exposed B103 cells, comparing with each sham exposure control (Fig. 3 B and C). However, when AKT activation was examined in 12 h-exposed HeLa cells and 24 h-exposed B103 cells, AKT phosphorylation levels remained constant in both the exposed and their sham controls (Fig. 3B, C). Considering the report that exposure to 60 Hz ELF-EMF at 0.8 mT activated NF-κB related signaling pathway in RAW264.7 cells [19], we also monitored the NF-κB activation in 60 Hz ELF-EMF-exposed HeLa (12 h) and B103 cells (24 h) at 14 mT and observed an elevation in the phosphorylation of the NF-κB p65 subunit (Fig. 3D, E).

**Figure 3.**
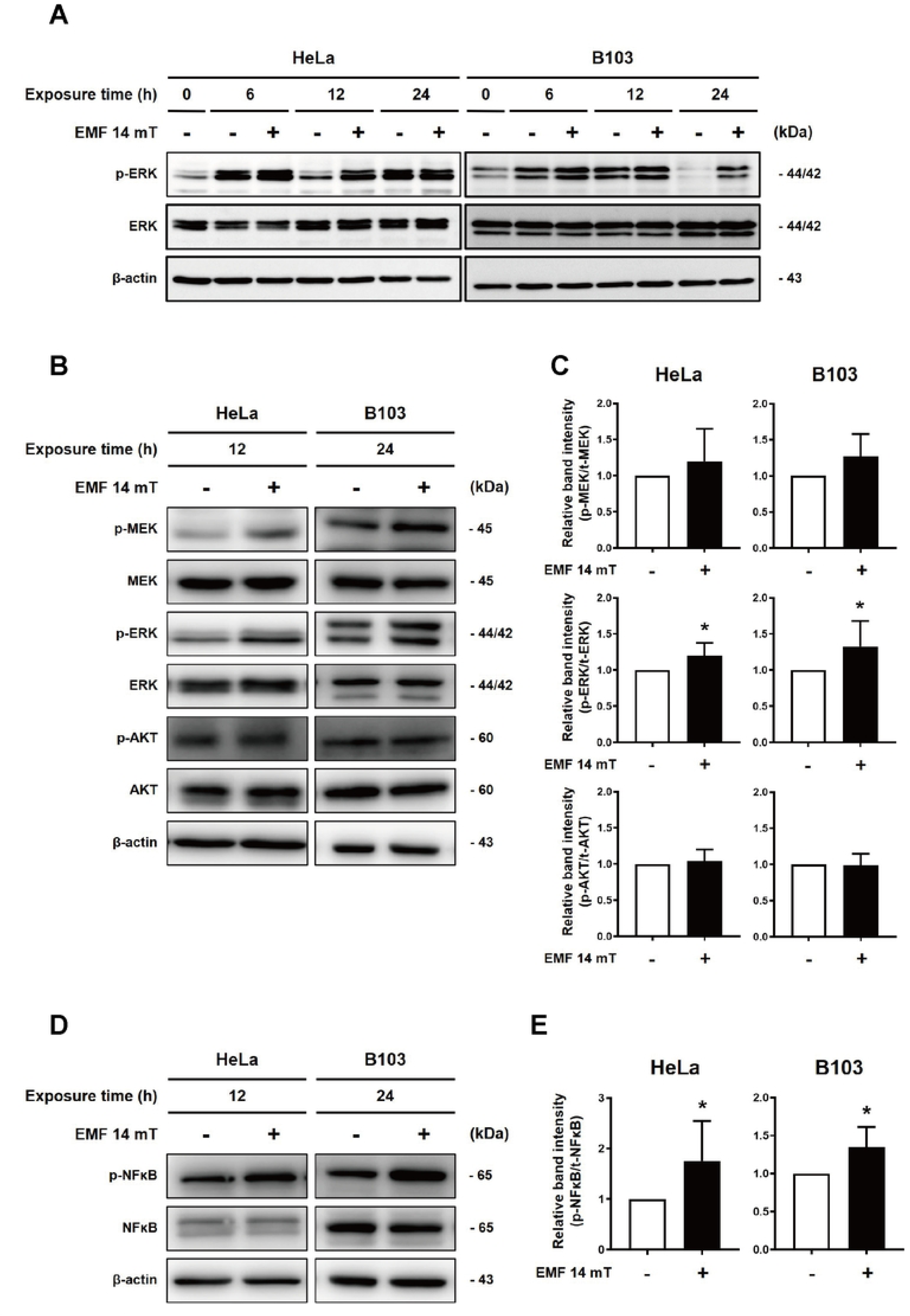
Continuous exposure to 60 Hz ELF-EMF at 14 mT increased phosphorylation of ERK1/2 and MEK1/2. HeLa and B103 cells were continuously exposed to 14 mT ELF-EMF for the indicated time period. (A) After exposure, the expression of p-ERK1/2 and ERK1/2 were immediately detected by western blot analysis. (B) The expression of p-MEK, MEK, p-ERK1/2, ERK1/2, p-AKT, and AKT was detected in HeLa and B103 cells by western blot analysis. (D) The expression of p-NFκB and NFκB were detected in HeLa and B103 cells by western blot analysis. (C, E) Bar graphs represent the relative expression of (C) p-MEK, p-ERK1/2, and p-AKT, and (E) p-NFκB over each sham exposure control in (B) and (D). Each band was normalized with actin and the relative band intensity of the phosphorylated form over unphosphorylated total was plotted. At least three independent experiments were performed and values are presented as the mean ± SD (n ≥ 3). P-values were determined by Maan-Whitney U-test and a value of *P<0.05 was considered statistically significant.

### Exposure to 60 Hz ELF-EMF induced slight modifications in the cell cycle profile

Considering a continuous exposure of 60 Hz ELF-EMF at 14 mT increased cell viability and ERK activation, we examined the DNA content of HeLa and B103 cells to confirm whether 60 Hz ELF-EMF exposure promotes cell cycle transition. HeLa and B103 cells were exposed to 60 Hz ELF-EMF at 14 mT for 6, 12, 24, 48, and 72 h respectively, and sorted by their DNA contents through flow cytometry. No significant changes in cell distributions by DNA contents at each time point were observed between the ELF-EMF-exposed and their sham controls (Fig. 4A). When we also checked the cell death of 60 Hz ELF-EMF-exposed cells at 14 mT for 72 h and the sham control by flow cytometry after FITC-conjugated Annexin V and PI staining, we could not detect any sign of cell death as expected (data not shown). Then, we examined the expression of cell cycle progression markers, cyclin-dependent kinase 4 (CDK4) for the G1 phase and proliferation cell nuclear antigen (PCNA) for the S phase by western blots, and no significant changes were observed in the cells exposed to ELF-EMF for 6, 12, and 24 h and their sham controls (Supplementary Figure S3).

**Figure 4.**
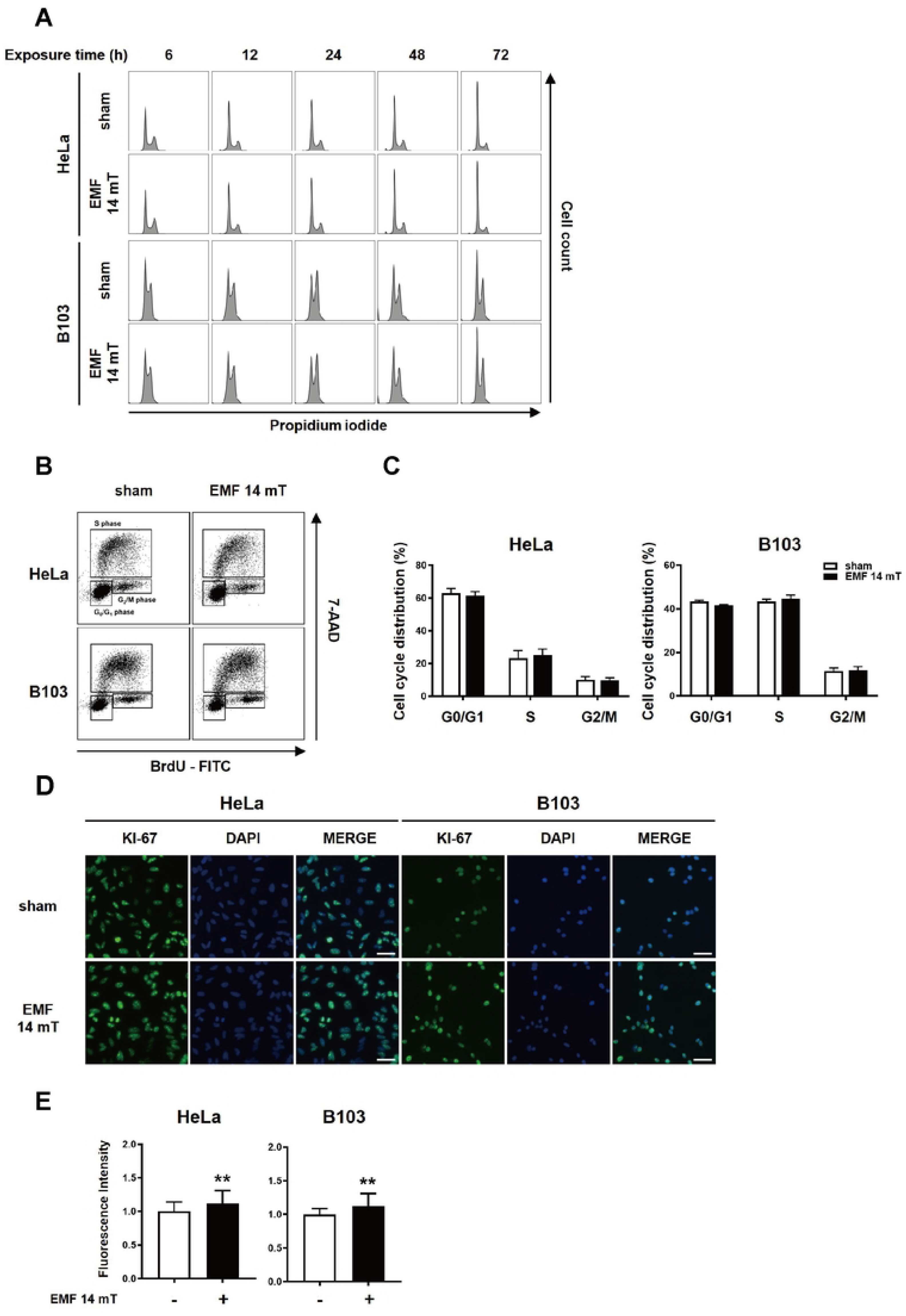
Continuous exposure to 60 Hz ELF-EMF at 14 mT induced slight modifications in the cell cycle profile. (A) Cells were exposed to 14 mT ELF-EMF for 6, 12, 24, 48, and 72 h. The DNA content of the ELF-EMF exposed and the sham control was analyzed by flow cytometry after PI staining. (B) BrdU incorporation and the DNA content of the 24 h-ELF-EMF-exposed cells and the sham control were analyzed by flow cytometry. (C) The distribution of cells in each cell cycle stage was calculated from (B) by FlowJo V10 software and plotted as the mean ± SD. (D) After exposure to ELF-EMF for 24 h, Ki-67 expression was detected with immunofluorescence analysis. (E) The relative fluorescence intensity of Ki-67 (the EMF-exposed/the sham control) in (D) was analyzed with ImageJ and plotted as the mean ± SD of more than three independent experiments. P-values were determined by Maan Whitney U-test and **P<0.01 was considered statistically significant.

For more accurate cell cycle analysis, HeLa and B103 cells treated with 60 Hz ELF-EMF at 14 mT for 24 h were probed with BrdU, and BrdU-incorporated cells were monitored by flow cytometry. While the S phase population increased slightly, the difference was not statistically significant, with a p-value of 0.1 (Fig. 4 B, C). To confirm the minor difference observed by BrdU incorporation, we assessed Ki-67 expression by immunofluorescence microscopy to define the proliferative status in HeLa and B103 cells continuously exposed to 60 Hz ELF-EMF at 14 mT for 24 h. Following ELF-EMF exposure, Ki-67 expression was increased both in HeLa and B103 cells, compared with the sham exposure control (Fig. 4 D, E). Altogether, these observations suggest that continuous exposure to 60 Hz ELF-EMF only slightly accelerates cell cycle progression for increased proliferation.

### The proliferative effect of continuous exposure to uniform 60 Hz ELF-EMF persisted in mitochondrial hypofunction

Several studies have reported that exposure to ELF-EMF generates reactive oxygen species (ROS) *in vivo* [18, 20–22]. ROS are consistently generated in cells by metabolic redox reactions. A minor increase in intracellular ROS levels may induce cell proliferation by activating signaling cascades [23]. Antioxidants also increase cell proliferation by reducing intracellular ROS levels [24, 25]. Thus, we questioned whether the increased proliferation by 60 Hz ELF-EMF was triggered by changes in intracellular and/or mitochondrial ROS levels.

First, we examined the intracellular ROS by measuring carboxyl-H2DCFDA in HeLa cells continuously exposed to 60 Hz ELF-EMF for 6 and 24 h at 14 mT. Human cells uptake non-fluorescent carboxyl-H2DCFDA, which is then oxidized by intracellular ROS to emit a bright green fluorescence signal. Immediately after the exposure, cells were incubated with 25 μM non-fluorescent carboxyl-H2DCFDA in the dark for 30 min and the intensity of fluorescence in these cells was monitored by flow cytometry. However, we could not detect any significant difference in fluorescent intensity between the ELF-EMF-exposed cells and the sham controls (Fig. 5A). Since the majority of ROS within cells originate from mitochondria, we then examined the mitochondrial ROS in HeLa cells continuously exposed to 60 Hz ELF-EMF for 6 and 24 h at 14 mT. Immediately taken from the exposure device, cells were incubated with 5 μM MitoSox for 30 min at 37°C and the intensity of the MitoSox fluorescent signal, which responds to mitochondrial superoxide, in these cells was monitored by flow cytometry. We could not observe any difference in the intensity of the MitoSOX fluorescent signal between the ELF-EMF-exposed and the sham control (Fig. 5B).

**Figure 5.**
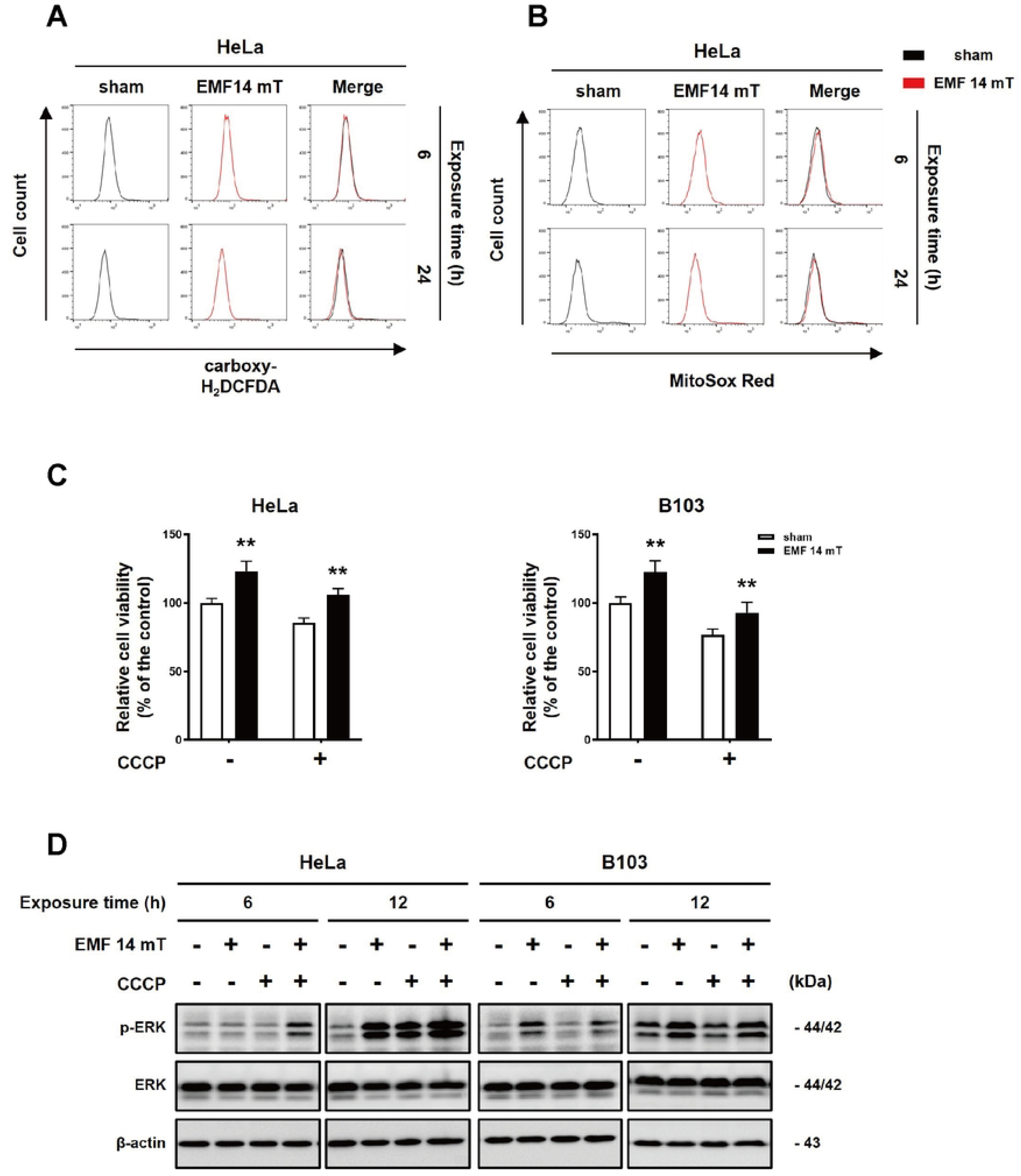
Continuous exposure to 14 mT 60 Hz ELF-EMF does not change intracellular ROS levels. (A, B) HeLa cells were exposed to 14 mT ELF-EMF for 6 h and 24 h. Intracellular ROS was probed with carboxy-H_2_DCFDA (A) and mitochondrial ROS was probed with MitoSox Red (B). Fluorescence of each cell was detected by flow cytometry. For each flow cytometric analysis, 10,000 cells were counted and plotted. The data are shown as overlapping histogram plots. (C) HeLa and B103 cells in the medium containing 3 μM CCCP (CCCP dissolved in DMSO) were exposed to 14 mT ELF-EMF for 72 h. After ELF-EMF exposure, cell viability was analyzed by MTT assay. In both CCCP-treated and -untreated (DMSO only) groups, the relative cell viability of ELF-EMF-exposed cells was plotted as a percentage of their sham control cells. Values are presented as the mean ± SD by three independent experiments (n=3) and P-values were determined by two-way ANOVA with multiple comparisons test. **P<0.01 was considered statistically significant. (D) HeLa and B103 cells in the medium containing 3 μM CCCP were exposed to 14 mT ELF-EMF for 6 and 12 h, and the expression level of p-ERK1/2 and ERK1/2 was detected by western blot analysis in both CCCP-treated and -untreated (DMSO only) cells with their sham exposure groups. ꞵ-actin from the same blot served as a loading control.

To further validate the effects of 60 Hz ELF-EMF on ROS generated in mitochondria, we exposed HeLa and B103 cells to 60 Hz ELF-EMF in the presence of mitochondrial uncoupler carbonyl cyanide 3-chlorophenylhydrazone (CCCP). In detail, HeLa and B103 cells were seeded for 18 h and they were replaced with a medium containing 3 μM CCCP, before these cells were exposed to 14 mT 60 Hz ELF-EMF for 72 h. The cells only pre-treated with DMSO were CCCP-untreated control. In our test, CCCP dissolved in DMSO decreased cell viability by 80% of the CCCP-untreated cells. After CCCP-treated HeLa cells were exposed to 14 mT ELF-EMF for 72 h, we examined the relative cell viability of the CCCP-treated and -untreated group to their ELF-EMF sham control cells. The relative cell viability of the CCCP-treated and -untreated groups showed similar increases by ELF-EMF exposure, 24.7%, and 21.3%, respectively, although the relative viability of the CCCP-treated group was consistently decreased to ∼ 80% of CCCP-untreated group. B103 cells also showed similar results: the relative cell viability of both the CCCP-treated and -untreated groups increased by 22.6% and 21.2% respectively by ELF-EMF exposure (Fig. 5C).

To confirm that the increased proliferation in CCCP-treated cells was also induced by ERK activation, the CCCP-treated and -untreated groups of HeLa and B103 were exposed to 14 mT ELF-EMF for 6 and 12 h and compared the ERK1/2 phosphorylation with their ELF-EMF sham control cells incubated for the same duration. When the cells were exposed to ELF-EMF, the increased expression of phospho-ERK1/2 was consistently observed both in the CCCP-treated and -untreated groups (Fig. 5D). Altogether, these observations strongly suggest that intracellular ROS are not responsible for the activated proliferation of 14 mT ELF-EMF-exposed human and rat cells through ERK activation.

### Exposure to 60 Hz ELF-EMF did not affect intracellular calcium level

To investigate the mechanism for ERK activation by 14 mT ELF-EMF exposure, we conducted RNA sequencing of 14 mT 60 Hz ELF-EMF-exposed B103 cells. We chose B103 cells for RNA sequencing because this cell type showed the most proliferative effect by ELF-EMF among the cell types used in this study. Unfortunately, we found that the changes in gene expression patterns between the ELF-EMF-exposed cells and the unexposed control were not substantial. Thus, we set a fold change threshold of 1.3 to detect differentially expressed genes (DEGs) more sensitively (Supplementary Fig. S4 A). Upon analysis based on the Kyoto Encyclopedia of Genes and Genomes (KEGG) pathway database, the calcium signaling pathways were considered the most likely candidates for the minute difference of gene expression pattern in ELF-EMF-exposed cells (Supplementary Fig. S4 B). Thus, we examined whether the calcium signaling pathways are responsible for ERK activation and increased proliferation in ELF-EMF-exposed cells. We measured the intracellular calcium level of HeLa and B103 cells with Fluo-4-AM after the cells were exposed to 14 mT ELF-EMF for 6 h and 24 h. However, there were no significant changes in basal calcium levels in the ELF-EMF-exposed cells, compared to their unexposed controls (Fig. 6A, B).

**Figure 6.**
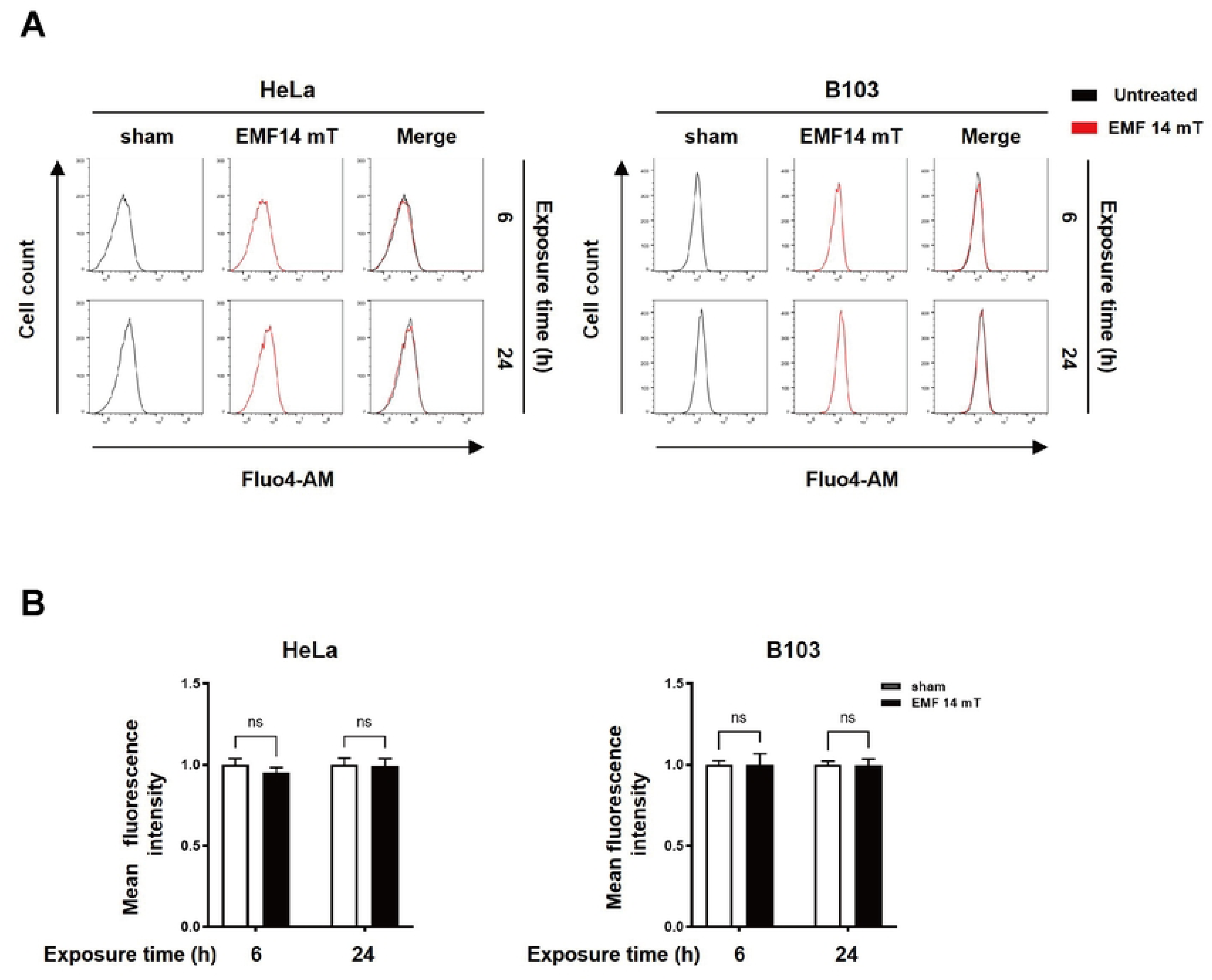
Exposure to 14 mT 60 Hz ELF-EMF had no effect on the basal levels of intracellular calcium. HeLa and B103 cells were exposed to 14 mT ELF-EMF for 6 h and 24 h with the unexposed sham control. Trypsinized cells were stained with Fluo4-AM and analyzed by flow cytometry. For each flow cytometric analysis, 10,000 cells were counted and plotted. The data are shown as overlapping histogram plots. (A) The x-axis represents fluorescence intensity indicating Fluo4-AM binding, and the y-axis shows cell count. (B) Mean fluorescence intensity (MFI) was calculated and plotted as the mean ± SD of three independent experiments and P-values were determined by Maan-Whitney U-test. ns means non-significant.

To further verify that the changes in intracellular calcium were not involved in ERK activation and increased proliferation by ELF-EMF exposures, we exposed HeLa cells to 14 mT ELF-EMF in the presence of a calcium chelator, BAPTA-AM. HeLa cells seeded for 18 h were treated with 10 μM BAPTA-AM and exposed to 14 mT ELF-EMF for 72 h to examine the relative cell viability of the BAPTA-AM-treated and -untreated groups to their ELF-EMF sham control cells. The relative viability of the BAPTA-AM-treated group was decreased to about 50% of the BAPTA-AM-untreated group because of the adverse effect of calcium chelation (Supplementary Fig. S4 C). However, compared to their ELF-EMF sham controls, the relative cell viability of the BAPTA-AM-treated and -untreated groups showed obvious increases by ELF-EMF exposure, 17.89%, and 45.41%, respectively (Supplementary Fig. S4 C). These observations strongly suggest that the changes in intracellular calcium level are not responsible for the activated cell proliferation by 14 mT ELF-EMF.

## Discussion

Most physiological effects of 50 ∼ 60 Hz ELF-EMF were reported at 1 mT or under, but we wanted to examine the effect of ELF-EMF at over 10 mT, which occurs around social infrastructures such as large power plants, factories, and telecommunication facilities. In this study, we designed a novel device to generate uniform 60 Hz ELF-EMF at 10 to 20 mT without producing heat. We then monitored the internal temperature of the device in the incubator at the magnetic flux density of 10, 14, 16, 18, and 20 mT and found a meaningful temperature rise at 18 and 20 mT (Supplementary Fig. S2). Thus, we investigated the cellular effects of 60 Hz ELF-EMF at 10 to 16 mT on various human cells. Since we increased the magnetic flux density of ELF-EMF to 10 mT or higher, we could not predict whether ELF-EMF induces pro-proliferative effects by activating signaling pathways or anti-proliferative effects by triggering DNA damage and cell death. We could not detect any sign of cell death or anti-proliferative effect by ELF-EMF exposure (data not shown). Rather, as shown in Figure 2, continuous exposure of 60 Hz ELF-EMF activated cell proliferation at 10 and 14 mT in all cell types proportionally to the strength of the magnetic flux density applied. At 16 mT, however, there was no further increase in cell proliferation as compared to that at 14 mT, and there was even a decrease in proliferation in IMR90 and Hep3B cells. These observations suggest that the proliferation-activating effect of ELF-EMF varies with the specific strength of the electromagnetic field and different cell types. Thus, we focused our study on the effect of 14 mT ELF-EMF, which activates proliferation in various rat and human cell types. In all cells whose proliferation was activated by continuous ELF-EMF exposure at 10 and 14 mT, we observed the activation of stress-activated MEK/ERK signaling pathways and NF-κB, but not of AKT.

Considering that continuous exposure to ELF-EMF at 14 mT for 72 h increased proliferation of various cell types by ∼20%, we examined the cell cycle of ELF-EMF-exposed cells. Although the cell cycle progression of G1 to the S phase was not significantly accelerated, there was a modest increase in S phase cells. Due to the minimal effect of ELF-EMF on cells, significant changes in cell cycle analysis were difficult to observe. Thus, we attempted to examine the proliferative state in the ELF-EMF-exposed cells. We observed an increase in Ki-67 expression, indicating more cells were in a proliferative state when exposed to ELF-EMF at 14 mT. We assume that 10-14 mT ELF-EMF exposure marginally activates the G1 to S transition in each cell cycle to be hardly detectable by cell cycle analysis, but results in the overall increase of proliferation by ∼20% after proceeding several cell cycles for 72 h exposure.

We observed promoted proliferation by ELF-EMF at 10-14 mT in both normal and cancer cells and proliferation was more increased in fast-growing cancer cells than in relatively slow-growing normal cells. These observations may support some epidemiological studies that suggest ELF-EMFs may accelerate tumor formation [7]. In other words, ELF-EMF exposure does not induce DNA damage and cytotoxicity to switch normal cells to fast-growing cancer cells, but it may accelerate tumor formation by promoting the proliferation of already mutated fast-growing cancer cells.

In order to understand the mechanism of ELF-EMF to promote cell proliferation by ERK activation, we first examined the changes in intracellular and mitochondrial levels of ROS, which are frequently highlighted in other ELF-EMF-related studies [11]. However, upon continuous exposure to 14 mT ELF-EMF for 72 h in our study, no changes in ROS levels were observed. Using mitochondrial uncoupler, CCCP, we further confirmed that intracellular ROS generated in mitochondria is not responsible for increased proliferation and ERK activation (Fig. 5 C and D). Thus, we concluded that the increased proliferation by 10-14 mT ELF-EMF cannot be explained by the fine-tuned regulation of intracellular ROS levels reported for the physiological outcomes by other studies [26. 27].

To obtain a clue for ERK activation and accelerated proliferation by ELF-EMF, RNA-seq transcriptomic analysis was performed with 14 mT ELF-EMF-exposed B103 cells whose proliferation was most highly increased among the cell types examined. However, as expected from the marginal change in cell cycle analysis, we could not detect differentially expressed genes with a fold change threshold of 2. When we decreased the threshold to 1.3, the KEGG pathway database suggested calcium signaling pathways to mediate the cellular responses by ELF-EMF exposure. However, when we examined the changes in intracellular calcium levels and the effect of a calcium chelator in the 14 mT ELF-EMF exposed cells, we could not detect any effect of calcium in the activation of cell proliferation and ERK activation.

In the present study, we tried to resolve plausible mechanisms including ROS and calcium to understand how exposure to 10-14 mT ELF-EMF promotes cell proliferation by activating ERK, but its mechanism is still an enigma. Considering that cell cycle and physiological changes caused by 10-14 mT ELF-EMF exposure are minimal, more delicate tools such as measuring minute fluctuation of plasma membrane potential or degree of the signaling receptor activation in a single cell level should be developed to differentiate the cellular variation induced by ELF-EMF exposure.

## Supporting information

**Supplementary Figure 1. Comparison of the overall structure between the previous and the revised ELF-EMF generating device**

A schematic front view of each device. The size and design detail of the previous and the revised model.

**Supplementary Figure 2. The monitored temperature value of inside and outside of the revised ELF-EMF-generating device**

A thermometer consisting of two probes was used to measure the temperature of the device and incubator simultaneously. Data were plotted as the mean ± SD (n ≥ 3). P < 0.01 (**) was considered statistically significant, and P > 0.05 was considered statistically non-significant (ns).

**Supplementary Figure 3. Continuous exposure to the 14 mT ELF-EMF does not affect the expression level of cell cycle marker proteins**

After exposure to 14 mT ELF-EMF for 6, 12, and 24 h, the expression of cell cycle stage markers was detected by western blot analysis with anti-Cdk4 and anti-PCNA. ꞵ-actin served as a loading control.

**Supplementary Figure 4. Transcriptome analysis in the ELF-EMF-exposed and - unexposed cells**

B103 cells were exposed to ELF-EMF for 24 h and mRNA expression was analyzed. (A) volcano plot of differentially expressed genes (DEGs) for ELF-EMF-treated cells vs - untreated cells. (B) A bubble chart of KEGG pathway enrichment analysis. The X-axis represents fold enrichment, and the Y-axis represents the pathway. The size of the bubble indicates the number of differentially expressed genes enriched in the pathway, and the color represents the significance. Fold enrichment indicates the proportion of differentially expressed genes (DEGs) in a specific KEGG classification compared to the total number of identified genes in that category. (C) HeLa cells were pre-treated with 10 μM BAPTA-AM and exposed to 14 mT ELF-EMF for 72 h. After the exposure, viable cells were analyzed by MTT assay and values are presented as the mean ± SD of more than three independent experiments. P-values were determined by two-way ANOVA with multiple comparisons test. **P<0.01 was considered statistically significant.

